# Predictive simulation of human movement in OpenSim using floating-base task space control

**DOI:** 10.1101/2024.02.13.580044

**Authors:** Nathaniel T. Pickle, Aravind Sundararajan

**Affiliations:** Biomedical, Energy and Materials Division, CFD Research Corporation, Huntsville, Alabama, United States of America

## Abstract

Task space control, also known as operational space control, is a useful paradigm for investigating neural control of human movement using predictive simulations. While some efforts have been made to implement task space control in the widely used open-source platform OpenSim, existing implementations do not support floating base kinematics, which is necessary for simulating gait and other types of human movement. Our aim in this work is to fill that gap. In this paper, we describe the theory and implementation of a floating base kinematics task space framework for torque- and muscle-driven simulations in OpenSim. Our framework builds on previous work that was limited to models with a base (i.e., root) segment fixed to ground. In addition, we integrate various algorithms from robotics in order to handle dynamically changing contacts and task prioritization. The framework can be used to generate realistic walking gaits by prescribing a small set of controller gains and gait parameters such as step length, step width and center of mass velocity. Task can be specified as desired positions, rotations, or higher-order feature such as base of support and whole-body angular momentum. We provide several examples to demonstrate how framework is successful in orchestrating a complex hierarchy of tasks that work in concert to perform both balance control and gait generation. The implementation is freely available for roboticists and biomechanists to use with OpenSim.

**Author summary:** Recent advances in computational biomechanics have provided researchers with tools capable of predicting human movement. Previous approaches to simulating human movement required experimental data as input and the simulation would replicate the experimental motion. This conventional approach limited the scientific insights to the specific movement recorded in the laboratory. With predictive approaches, researchers can investigate how a person might respond to various factors, such as reduced muscle strength or an assistive device such as a robotic exoskeleton. Existing approaches for generating predictive simulations utilize optimization-based approaches, which can be time-consuming and difficult to troubleshoot. Task space control is an alternative approach which is widely used in robotics. Conceptually, task space control aims to generate a simulation by specifying “tasks”, such as moving a hand or foot to a desired position, and computing the joint angles required to achieve the task. Here we aim to outline the mathematics behind task space control and demonstrate how task space control can be used to generate simulations of movements such as walking.

## Introduction

Predictive simulations are a major emerging advance in the field of computational biomechanics. Conventional simulation techniques have utilized a tracking approach, which requires experimental motion capture and force data as input. Simulations are generated by tracking the recorded joint angles. A commonly used method is computed muscle control (CMC), in which desired joint torques are computed using an inverse dynamics approach, then muscle forces are computed using an optimization procedure that assumes the primary neural objective is to minimize metabolic energy expenditure, represented mathematically as minimizing the sum of squared muscle activations [1]. Tracking simulation paradigms such as CMC have several inherent limitations. First, while they are useful for investigating muscle forces and coordination during a measured motion, tracking simulations cannot predict response to a perturbation. As a consequence, human responses to external perturbations such as tripping over an obstacle, receiving assistance from a device, carrying a heavy load, or experiencing muscle fatigue cannot be simulated in the absence of experimental data. A related consequence is that tracking simulations are often “brittle”. Errors and assumptions in model parameters such as body segment inertia properties and joint locations introduce inaccuracies that can make it difficult or impossible for a numerical solver to find a valid solution, a problem known as dynamic inconsistency [2]. To address this problem, researchers utilize approaches that make small adjustments to the model mass and kinematics [3]. These adjustments improve dynamic consistency in the simulation, but frequently at the expense of realism of the simulation. For example, the model may drift vertically or take a step that no longer aligns with recorded force data. Predictive simulations achieve a higher level of robustness by prioritizing physical realism over exact replication of experimental kinematics. Thus, a predictive simulation may not exactly replicate a specific measured motion, but will respect realistic physics such as ground contact and controlling balance. Furthermore, because predictive simulations utilize more abstract goals or objectives (e.g., foot trajectory or keeping the center of mass within the base of support) and compute joint angles required to achieve those objectives, predictive simulation paradigms offer the potential to gain novel insights into underlying neural objectives that govern human movement.

Predictive simulation paradigms have been utilized by a number of biomechanists. One approach is direct shooting, in which an optimization routine in which sets of input parameters are sought that achieve some optimization criterion, such as maximizing distance traveled [4]. Other approaches utilize evolutionary algorithms, such as the Covariance Matrix Adaptation Evolutionary Strategy (CMA-ES) implemented in the software tool SCONE [5]. Another widely used approach is direct collocation [6], [7], [8]. In direct collocation, the system dynamics are transcribed to a set of algebraic equations that can be solved using an optimization procedure. The OpenSim Moco tool [7] provides a software interface for direct collocation in biomechanical simulations. An approach that has received less attention from the biomechanics community is task space control (TSC).

The key feature of TSC is that desired movements are defined as goals within Cartesian spatial coordinates (3 orientations and 3 positions from either global reference frame or an inertial reference frame on a rigid body). These goals are then projected into joint space using the system Jacobian, which is a variational map relating spatial velocities of rigid bodies to generalized velocities of degrees of freedom (DOFs) in the model. TSC is a commonly used approach in real-time control of legged robots [9], [10] and animated characters [11], [12]. In addition, tasks are a natural way to represent aspects of the locomotor repertoire of humans and other animals. Examples of these features or tasks that are straightforward to express in the task space include managing the positions of the feet, the orientation of the trunk, and the position of the center of mass (COM) in order to produce stable walking gaits.

TSC is an attractive alternative framework for generating predictive simulations because it is possible to implement in an on-line paradigm (assuming a model of appropriate complexity). In contrast, direct shooting requires many iterations to optimize the controller and direct collocation requires the entire time history of the states to be converted to a large optimization problem. TSC may enable novel applications in real-time control of devices (e.g., prostheses and exoskeletons) or long-duration simulations. Some efforts have been made to make TSC available to biomechanists [13], [14], [15] by implementing it in the open source musculoskeletal simulation platform OpenSim [16]. However, these existing implementations have the significant limitation that they do not support floating base kinematics, where the root segment (commonly the pelvis) is unactuated and free to move relative to the world reference frame [10], [17]. Existing implementations can be used to simulate tasks such as reaching or cycling, where the pelvis is fixed, but are not suitable for generating simulations of walking, running, jumping, or other common movements.

Our objective in this work was to build upon the prior efforts of other researchers to implement a TSC framework in OpenSim that supports floating base kinematics, enabling predictive simulation of gait and other human movements. In addition, we outline approaches to resolving practical issues that arise such as avoiding joint limits and singularities. Our ultimate aim is to facilitate use of TSC in the biomechanics community so that researchers can more easily leverage it to explore the principles governing neural control of movement. In this paper, we provide a mathematical overview of floating base TSC. We demonstrate how our implementation can be used to generate predictive movement simulations in OpenSim.

## Materials and methods

First, we rigorously formalize the TSC framework by bringing together various previously separate notions of projection within subsequent manifolds that are used in robotic control: underactuation, task prioritization and hierarchy, as well as constraint or support consistency. For detailed mathematical proofs of the concepts presented here, we refer the reader to excellent publications by Stanev and Moustakas [14], [18] and Mistry and Schaal [17]. Our objective is to provide a rigorous yet accessible explanation of the concepts involved in TSC that provides biomechanists with a sufficient grasp of the method to be able to use it to formulate novel research questions in neuromechanical coordination of human movement.

As a preliminary, for any given variable *x*, we will use the notation 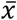to represent a dynamically consistent quantity (adheres to system constraints) and *x*^*∗*^to represent a matrix pseudoinverse. Dot notation 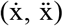 represents derivatives with respect to time unless otherwise stated. Dimensionality is represented by *n* for generalized coordinates, *c* for constraints, and *d* for tasks. Matrices or other quantities which are a function of the generalized coordinates *q* and their derivatives 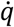,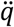will be indicated as such on the first usage, such as *M(q)*, but subsequently the arguments will be omitted for brevity.

### Equations of motion in configuration space and task space

The key concept in formulation of TSC is the distinction between task (or operational) space and configuration (or joint) space. It is common in both robotics and biomechanics to express the equations of motion of the robot or human in configuration space, as it is a concise representation that takes advantage of the kinematic constraints on system motion that are provided by linkages between segments in the system. Expressing the configuration of a linked multibody system is also natural in joint space, compared to expressing translations and rotations of each body in Cartesian coordinates in the operational space. However, desired tasks for a robot or simulated human to perform are often more readily expressed in Cartesian space. For example, we may wish to specify a desired position for an end effector (e.g., foot, hand) or a desired force or torque applied by that end effector in Cartesian space.

In configuration space, the equations of motion for a system with *n*_*q*_ degrees of freedom (i.e., generalized coordinates) are given by

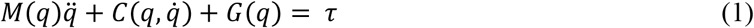

where 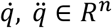 are the generalized positions, velocities, and accelerations, respectively, *M(q) ∈ R*^*n*×*n*^ is the system mass matrix, 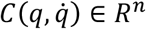 are the centrifugal and Coriolis contributions, *G*(*q) ∈ R*^*n*^ are the contributions of gravitational forces on each body represented in configuration space, and τ is the net generalized forces. It is worth noting that τ comprises contributions from various sources. We could instead express τ as

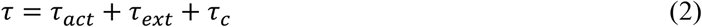

where τ_act_ are the actuator forces and moments, τ_ext_ are the contributions from external forces, and τ_c_ are the contributions from constraints. It is also worth noting that care must be taken with the sign conventions used in various derivations. OpenSim utilizes the Simbody physics engine, which uses the following sign convention:

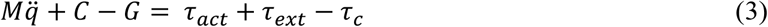

The mechanism for switching from configuration space to task space is the Jacobian [19]. The Jacobian is a matrix of first-order partial derivatives that relates changes in the generalized coordinates to corresponding changes in Cartesian coordinates at a specified point on the linked multibody system. We define the tasks positions, velocities, and accelerations as 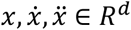, where *d* is the net dimensionality of the task (three for position or orientation tasks, one for a coordinate task). The Jacobian *J*_*t*_(*q) ∈ R*^*d*×*n*^ for a given task relates velocities in task space to velocities in configuration space,

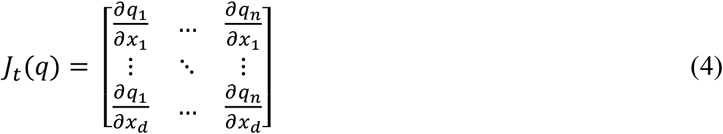

We can use the Jacobian to transform forces *F* expressed in task space to configuration space, mapping *F* ∈ *R*^*d*^ → τ ∈ *R*^*n*^:

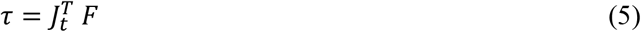

Using this relationship, the system equations of motion can be transformed from configuration space to task space, represented as

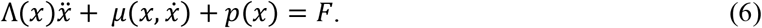

Here, 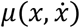 represents the centrifugal and Coriolis forces, and *p*(*x*) represents the gravitational forces, both in task space. The term Λ(*x*) ∈ *R*^*d*×*d*^ is the system inertia matrix in task space, and is derived as the *M(q)*-weighted pseudoinverse of *J*(*q):*

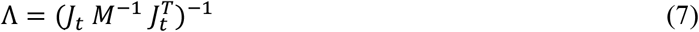

### Tasks

With the mathematical foundation in place, we can now begin to define desired movements in task space. In this TSC framework, we derived three essential categories of tasks (Table 1). These tasks are defined using OpenSim’s nomenclature. An arbitrary number of tasks can be added to any simulation, though computational performance will decline as the number of tasks grows.

**Table 1:** Types of tasks available in the OpenSim implementation of task space control.

| Task | Description |
| --- | --- |
| **Station** | 3-dimensional task that drives a specified "station" point on a rigid body to a desired position in 3-dimensional space. |
| **Orientation** | 3-dimensional task that drives a specified “frame” on a rigid body to a desired orientation in 3-dimensional space. |
| **Coordinate** | 1-dimensional task that drives a specified coordinate to a desired value. |

Station and Orientation tasks specify a point or orientation, respectively, in Cartesian space. Coordinate tasks are expressed in configuration space rather than task space, but are included due to their utility in joint limit avoidance or generating a default “return to nominal configuration” behavior.

Each type of task can be used to compute desired accelerations in task space. Here we utilize a standard proportional-integral-derivative (PID) feedback control approach to compute the desired accelerations 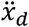 as

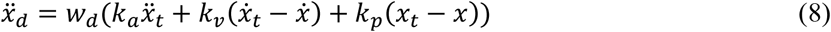

where *k*_*a*_, *k*_*v*_, and *k*_*p*_ are the acceleration, velocity, and position gains, respectively, 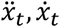 and *x*_*t*_ are the specified task acceleration, velocity, and position, respectively, and 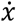 and *x* are the current velocity and position, respectively, in task space. We also apply a weight *w*_*d*_ which can be used to adjust the relative influence of a task. We can substitute 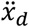 into Equation (6) to compute the desired forces in task space.

### Dynamic consistency with constraints

A commonly exploited characteristic of task space representations of movement is the ability to formulate joint-space movements which exist in the null space, or kernel, of system constraints. Intuitively, this means we can identify movements of the joints which produce no reactive force or moment at the constraint location. An example would be performing the task of turning an imaginary doorknob, where the hand position is fixed in space by a point constraint but is free to rotate. The hand remains at the same location in space while the rest of your arm rotates by moving the wrist, elbow, and shoulder joints. Null space projection can be used to compute actuation forces which are dynamically consistent with system constraints; that is, the actuation forces operate in the null space of the constraints.

We define a set of constraints on the system as

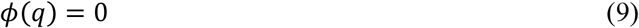

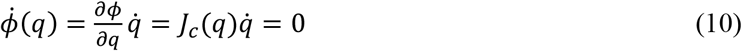

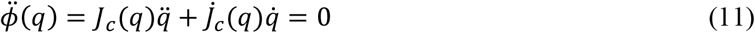

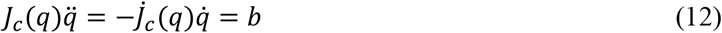

where *J*_*c*_(*q*) ∈ *R*^*c*×*n*^ is the constraint Jacobian. We can further define

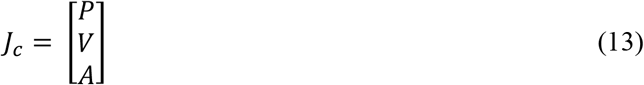

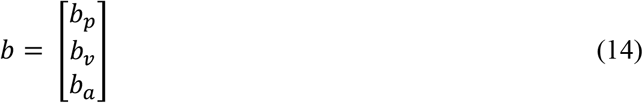

Here *P, V*, and *A* are the position, velocity, and acceleration components, respectively. The constraint vector *b* comprises *b*_*p*_, *b*_*v*_, and *b*_*a*_, which are a set of algebraic constraints on the position, velocity, and acceleration, respectively. The quantity 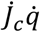 is also referred to as the constraint bias. The algebraic constraints must satisfy a set of equations that express the position, velocity, and acceleration level constraints [20].

Any subsequent tasks, in order to adhere to the constraint manifold, must operate within the kernel of this linear operator *J*_*c*_ ∈ *R*^*d*×*n*^ where *J*_*c*_^*∗*^ is a pseudoinverse of *J*_*c*_. Various pseudoinverse methods can be used, each with different properties. The Moore-Penrose pseudoinverse is a commonly used pseudoinverse, and is given by

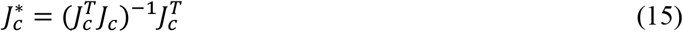

when *J*_*c*_ is full column rank (left inverse), such that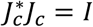. The Moore-Penrose pseudoinverse produces a least squares solution for dynamically consistent actuation forces and torques. Alternatively, the singular value decomposition (SVD) can be used to compute the inverse. In general, the singular value decomposition (SVD) of a given matrix *A* is given by

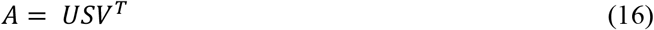

where the matrices *U, V* are orthogonal while *S* can be considered a projection operator that maps *U* → *V*. The pseudoinverse of *A* is given by

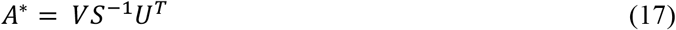

The constraint Jacobian *J*_*c*_ and its pseudoinverse can be used to compute an orthogonal projection operator *N*_*c*_ ∈ *R*^*n*×*n*^ that projects the actuation torques into the null space (i.e., kernel) of the constraints [18]. Because we typically have more DOFs than constraints (*n* > *c*), we utilize the projection onto the left null space, definitionally given by

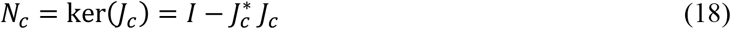

We can then filter the actuation torques through the constraint null space projection operator to produce the dynamically consistent joint torques 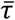 for a desired task force *F*:

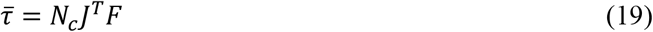

### Task prioritization

Nearly all human movements require multiple tasks to be defined in order to produce the desired motion. For example, gait requires control of the COM relative to the base of support, achieving a desired COM velocity, controlling trunk posture, and moving the feet. Some tasks, such as maintaining balance to avoid falling, may take precedence over other tasks. In order to simulate this behavior, we require a mathematical framework to represent task prioritization, where the achievement of high priority tasks must not be disrupted by lower priority tasks.

Similar to the approach outlined in Section 0 for constraints, task prioritization can be accomplished by filtering lower priority tasks through the null space of higher priority tasks. In terms of implementation, we choose to create stacked Jacobian matrices representing the tasks at each priority level [17] rather than a recursive algorithm [14].

At each priority level *i*, we will have a total of *k* tasks, and we create an augmented Jacobian matrix *J*_*a*|*i*_ by stacking the tasks at priority level *i*:

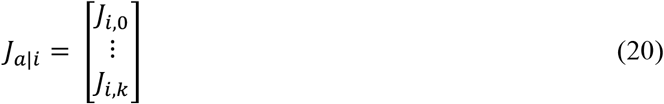

Note that at each priority level we append new task Jacobian matrices to *J*_*a*|*i*−1_, so the augmented Jacobian grows at each subsequent priority level. We use the augmented Jacobian to form the null space projection matrix for tasks at priority level *i*:

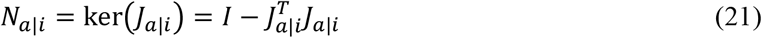

We can then recursively calculate the control torques for each priority level, filtering them through the null space of higher priority tasks in order to prevent interfering with those higher priority tasks:

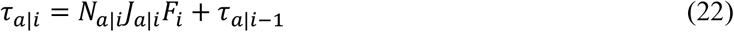

Tasks must never interfere with holonomic constraints in the model, which can be considered a priority “-1” task. We use the total aggregate Jacobian *J*_*a*_of all tasks to identify the dynamically consistent generalized forces that satisfy the constraints of the aggregate task manifold (see Section 9). The minimum kinetic energy solution for τ_*a*_ is computed by modifying Equations (6) and (7) to project *M*^*−1*^ through the constraint null space *N*_*c*_ [10]. First, we compute the dynamically consistent mass matrix expressed in task space:

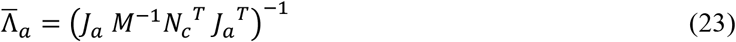

We perform the matrix inversion operation using a damped SVD pseudoinverse to avoid singularities (see Section 0. Next, we compute the dynamically consistent aggregate Jacobian 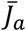 as

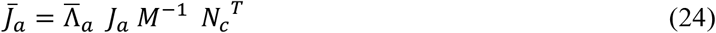

and construct the null space projection matrix

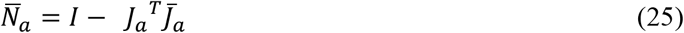

We can now compute aggregate control torques which adhere to our desired task prioritization as well as the holonomic constraints in the system:

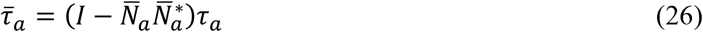

### Floating base kinematics

The derivations presented in the previous sections are sufficient only for generating simulations of systems in which the base segment (commonly the pelvis in biomechanical simulations) is fixed in space. For example, these equations can be used to simulate a reaching task where the torso is fixed, or cycling where the pelvis is fixed to the bicycle seat. The key contribution we present is to extend existing TSC implementations for biomechanical applications [13], [14], [15] in order to accommodate systems in which the base segment is free to move in 6DOF. To accomplish this, we refer to approaches developed by Mistry and Schaal [17] and Sentis [10].

In the case of a floating-base system, the base segment is free to move in 6DOF but is not directly actuated in any of those DOFs. However, that does not mean that the base DOFs are not controllable. Constraints and external forces applied at other segments, such as the end effectors, can be used to indirectly control the motion of the base segment. Conceptually, this is the opposite approach to that outlined in Section 0: rather than computing torques which exist in the null space of the constraints, we wish to intentionally leverage external forces and constraints to control the motion of the base segment as if it were actuated.

To represent the lack of actuation at the base segment, we construct a selection matrix *B* which acts as a constraint on the actuation torques:

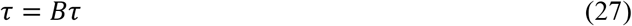

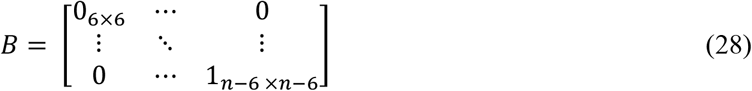

Note that we assume here that the (unactuated) base segment DOFs are represented by the first six rows. The selection matrix can also be used to indicate coordinates other than the base segment which are unactuated, but here we focus our discussion only on the base segment.

Often, operational space formulations of mobile base systems decouple the 6 virtual DOFs of the base from the robot coordinates and define task Jacobians from the base inertial frame; however, for simplicity, this is not done here, but consideration of the virtual DOFs is done internally by Simbody Engine when constructing frame or station task Jacobians.

In a system with floating base kinematics, the base segment is not directly actuated. However, we wish to develop an architecture that allows us to control these DOFs indirectly using interaction with the environment, which can be modeled either as kinematic constraints or as external forces. The base segment DOFs are treated as “virtual” DOFs, and the task space are filtered using an underactuation matrix *B* [17]. Using the dynamically consistent aggregate nullspace 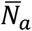 (Equation (25) and *B* (Equation (28).

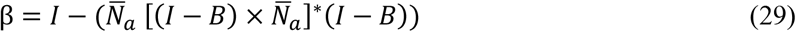

The aggregate generalized forces 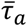 that satisfy the requirements of all tasks is then

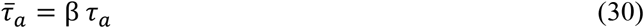

Note that in the case of full actuation or a model that has no virtual DOFs like a classical manipulator, *B* = *I*, so β simplifies to *I* and Equation (30) simplifies to Equation (26).

### Joint limits weighting matrix

Simulating human movement requires anatomically realistic limits to be placed on joint ranges of motion. Conventional approaches to enforcing joint limits include kinematic constraints and passive forces expressed as a function of joint angle [21]. However, those approaches each have limitations. Utilizing kinematic constraints introduces discontinuities into the system, and does not accurately represent the development of passive forces near the extremes of joint range of motion. Passive forces computed as a function of joint angle provide a more realistic representation of soft tissue mechanical behavior, but are not accounted for the by control algorithm.

For coordinate tasks, typically of the form, *J* ∈ *R*^1×*n*^ = [0 … *a* … 0], the constant *a* can be chosen such that the joint ranges of motion are not passed. While many methodologies for joint limit avoidance have been studied [22], [23], [24], the repulsive potential field by means of the Gradient Projection Method (Khatib 1986) was implemented in configuration space with a C1 continuous replacement for *a*, providing smooth transitions at the joint limits. For a 10% buffer *φ* from the maximal or minimal limit of a particular DOF, *a* is selected such that

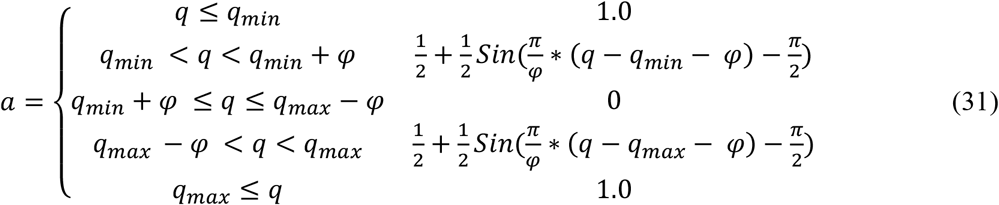

An example coordinate limit curve is shown in Figure 1.

**Figure 1:**
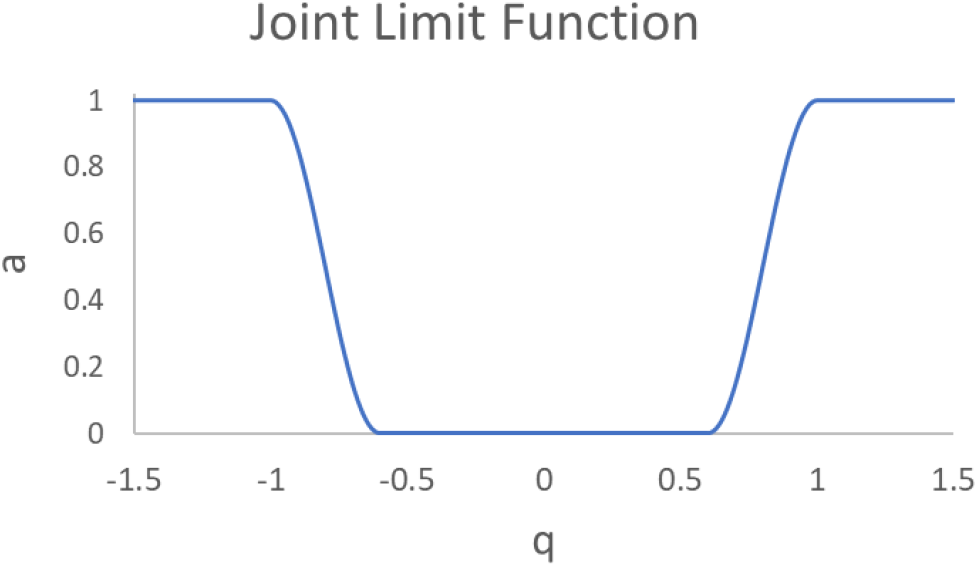
Illustration of the joint limit function *a* for joint angle *q* ranging from -1 to +1 radians with φ=0.4.

### Singularity avoidance

Kinematic singularities occur often, particularly in gait, where the task Jacobian becomes rank deficient in certain postures such as full extension of the knee or the alignment of various rotational axes in the arm. In singular configurations, the SVD (Equation (16) of the task Jacobian *J* reveals that the singular values matrix *S* has at least 1 zero value on the diagonal. Values sufficiently close to zero on the diagonal of *S* indicates that computing the inverse of *J* will cause loss of control or require infinite generalized accelerations to satisfy the demands of the task. We can mitigate these numerical instabilities in the SVD pseudoinverse calculation by adding a damping parameter ρ≪1 to the projection operator *S*:

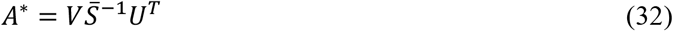

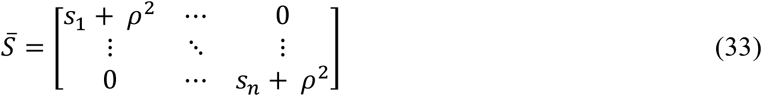

A selectively damped pseudoinverse can be implemented that damps the diagonal of *S* depending on a user-selected threshold based on the closeness of the singular value to 0. In this paper, we use the threshold 10^-15^, and also set ρ=10^-15^ if any *s*_*i*_ is below the threshold.

### Support-consistent matrix

Another important aspect required for generating simulations of gait and other human movements is the ability to represent contact with the environment, such as ground reaction forces at the feet. We have previously discussed a method for accommodating kinematic constraints (Section 9), which are commonly used to model ground (or other environmental) contact in robotics applications where the ground reaction forces are unknown. However, in biomechanical simulations we can take advantage of full knowledge of ground reaction forces by modeling contact between the foot and ground.

To incorporate contact forces into our controller, we again turn to the Jacobian. Here we compute the Jacobian for the point of application of the external forces. In the case of gait, the point of contact is the center of pressure (COP). In our implementation we compute the COP for each body in contact with the environment, as each body may have multiple contact objects attached. *J*_*s*_^*k*^ ∈ *R*^6×*n*^ is the frame (position and rotation) Jacobian to the center of pressure (COP) on the *k*^th^ body in contact with the environment. We form the net support-consistent Jacobian *J*_*s*_ by concatenating the support Jacobian matrices for each point in contact with the environment. Thus we have

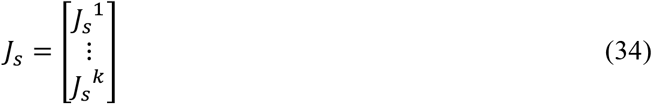

for *n*_*k*_ bodies in contact with the environment. We then utilize *J*_*s*_ in place of *J*_*c*_ in Equation (18). This is an approximation which effectively treats each external force as a weld constraint.

### Implementation in OpenSim

An important objective of this work was to implement the TSC framework natively in OpenSim, such that it utilizes existing OpenSim functionality and can be used similarly to any other control schema available in OpenSim. To achieve this goal, we relied heavily on prior efforts by Stanev and Moustakas [14] and the Stanford University development team [13], who made significant advances in implementing TSC in OpenSim and Simbody. Our key contributions were to extend the existing functionality by adding support for floating base systems and joint and singularity avoidance. In addition, the implementation utilizes OpenSim’s component architecture such that tasks can be serialized and deserialized in XML format.

### Gait controller example

The various aspects of the TSC framework described in the preceding sections provide the building blocks necessary for generating simulations of human movement. However, the next challenge is to utilize the components such that a realistic motion is generated. Tasks provide the mathematical machinery for tracking desired positions or orientations in task space, but we must still determine what those desired positions and orientations should be. To demonstrate how the TSC components can be utilized to simulate human movement, we present an example in which we generate a simple 2D simulation gait in a lower-body model of a human.

### Prioritization scheme

Bipedal gait can be generated from a set of high-level “features” [11], which are readily represented in task space. To achieve gait, we implemented a prioritized set of features (**Table *2***).

**Table 2:** Features of gait for natural walking. Priority 0 is the highest priority level.

| Priority Level | Task | Type |
| --- | --- | --- |
| **0** | Torso orientation | Orientation |
| **0** | Pelvis forward progression | Station |
| **0** | Pelvis height | Station |
| **0** | Pelvis orientation | Orientation |
| **1** | Swing foot position | Station |
| **1** | Swing foot orientation | Orientation |

### Foot position

The model will keep the stance foot stationary in order to maintain control of the upper body, so no tasks are necessary on the stance foot. The desired position for the swing foot needs to be specified as a trajectory from the current location to a new foot placement location. For this example, we use a simple sinusoidal trajectory, though more sophisticated foot placement algorithms could be substituted. We define the tracking function as

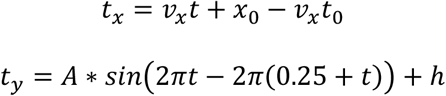

where *t*_*x*_ and *t*_*y*_ are the desired horizontal and vertical foot positions, respectively, *t* is the current time, *v*_*x*_ is the desired foot horizontal velocity, *x*_*0*_ is the initial foot horizontal position at the initiation of swing, and *t*_*0*_ is the time at initiation of swing. For the vertical position, *A* is the amplitude of the sinusoid and *h* is an offset applied to account for tracking error in the foot position.

### Posture: pelvis and torso control

Three tasks are used to control overall posture of the model during gait. The pelvis position is used as a surrogate for the COM, and is maintained at a constant height equal to the initial height. This is a simplification, as in actual human gait the COM motion can be described by an inverted pendulum model [25]. Pelvis forward progression is controlled by commanding the model to maintain a constant distance in front of the center of the base of support, which is computed as the midpoint between the calcaneus bodies in the anterior/posterior direction. The torso orientation is maintained constant at the initial orientation to maintain an upright posture.

### State machine for gait generation

The gait controller must adjust the desired center of mass and foot positions, i.e. controlling the upper body and swing foot while keeping the stance foot planted. Here, we developed a state machine that defines the swing task position as a function of time and the expressed body, after the foot reaches the desired position, the functions are updated and applied to the contralateral foot.

## Results

Results for the simulated gait controller are shown in Figure 2. Large spikes in the reaction forces were observed at foot contact (Figure 2C), which also produced rapid changes in joint angles (Figure 2B) which are uncharacteristic of actual human gait.

**Figure 2:**
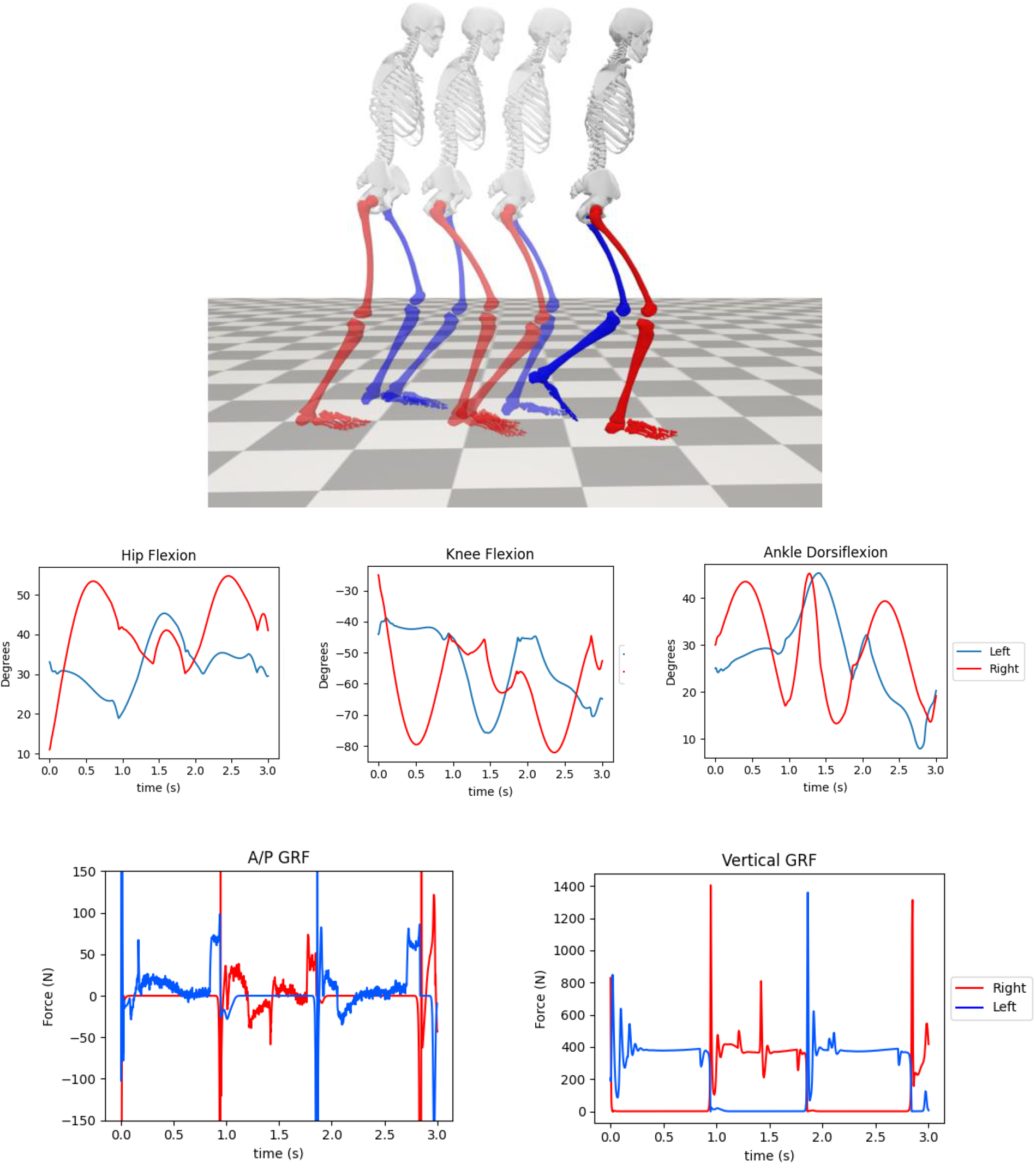
Gait controller results for the left leg (blue) and right leg (red). (A) Images illustrating the progression of two complete gait cycles, from stance to stance of the same leg. (B) Kinematics of the right leg over the course of the stride. (C) Ground reaction force patterns.

## Discussion

Task space control is a paradigm for generating predictive simulations of human movement that has significant potential to uncover novel insights into neural coordination. The objective of the work presented in this paper was to make key extensions to existing TSC implementations to facilitate simulation of gait, make the TSC framework readily available within the OpenSim musculoskeletal simulation software, and provide a conceptual overview of TSC for biomechanists interested in using TSC in their research. The task space control code is available as part of OpenSim Core (https://github.com/opensim-org/opensim-core) and examples are provided for C++.

Predictive simulation paradigms in computational biomechanics have recently emerged as a major development in the field. Whereas conventional simulation approaches require experimental data as input, predictive methods can generate human movement *de novo*. Direct collocation in particular has become a popular approach for generating predictive simulations. While direct collocation has been used in biomechanical simulations for a number of years [6], [26], [27], use of direct collocation in research initially required deep expertise in both the mathematics of direct collocation as well as the requisite programming skills to implement the calculations. After the release of the Moco tool in OpenSim [7], the popularity of direct collocation has grown tremendously. At the time of this writing, a Google Scholar search for the terms “OpenSim Moco” returns 47 results for the year 2022 and 65 results for the year 2023. The release of a user-friendly tool has substantially widened the pool of researchers who are able to utilize direct collocation in their research. Likewise, the SCONE tool has facilitated adoption of CMA-ES through release of a user-friendly software application. SCONE offers a useful alternative to Moco through features such as the ability to perform closed-loop control [28].

Our objective in implementing TSC in OpenSim is to provide researchers in biomechanics with another tool in their predictive simulation toolbox, which can serve as a complement to other tools such as Moco or SCONE. Compared to Moco, TSC may require less computational time to perform a simulation. Moco utilizes an optimization procedure to compute optimal controls and states, which typically requires hundreds or thousands of iterations to reach convergence. In contrast, TSC uses linear algebra to directly compute the control torques at each timestep (for a joint-driven simulation). Performing a muscle-driven simulation with TSC is still computationally intensive, as the muscle forces are computed using an optimization scheme as in OpenSim’s CMC algorithm. Another advantage of TSC is that the contribution of individual tasks and other terms can be queried during the simulation to aid in troubleshooting. Direct collocation requires the equations of motion to be transcribed to a set of algebraic constraints, at which point the optimizer takes over and attempts to find a solution. The optimization process can often be difficult to troubleshoot, as the root cause of failed simulations can be challenging to identify. Because TSC sequentially solves for the controls at each timestep, researchers may be better able to identify issues such as saturated actuators that may contribute to a failed simulation. In addition, in the future warnings can be added to identify infeasible tasks, a common source of failure in a TSC simulation. Another advantage of TSC is that because it solves for the controls sequentially, it is feasible to generate simulations of arbitrary duration. A simulation of an hour-long hike through mountainous terrain could theoretically be performed using TSC, and could even be run in real-time with appropriate modeling choices. In contrast, Moco solves for the entire motion simultaneously, which inherently limits the duration of movements that can be simulated and precludes the use of Moco for real-time simulation. However, this is also a limitation of TSC, as the controller does not utilize a look-ahead window to compute optimal controls accounting for the entirety of the motion. Moco and TSC are best understood as different tools for solving similar problems, and the appropriate choice of tool depends on the research question being asked.

A simple gait controller was presented to demonstrate how the TSC framework might be used to develop predictive simulations of human movement. The results presented here do not replicate the salient characteristics of human gait with sufficient accuracy for studying human gait. The simple task hierarchy and state machine produced joint kinematics and GRFs which only approximated actual human gait. Definition of a minimal set of high-level features that produce realistic gait is an area of active research [29]. Issues such as stability of the feet during stance, correct selection of contact parameters, and more nuanced task definition and hierarchy are needed. However, the preliminary results presented here provide a foundation on which other researchers can build. The robotics literature has an extensive and rich body of knowledge regarding the development of gait controllers for bipedal locomotion. Application of those algorithms to the OpenSim TSC framework has the potential to yield exciting new insights into neural control of human movement.

The implementation of TSC described here has limitations which should be noted. Currently the TSC framework is best suited for real-time control of torque-driven models. As mentioned previously, the key challenge of muscle-driven simulation (in both TSC and CMC) remains the muscle redundancy problem. The majority of the computational time at each discrete timestep is spent solving for muscle controls that satisfy the demands of the task using either nonlinear optimization or computational geometry for prescribed kinematics. Future advances in computational speed or borrowed techniques from Kalman filtering, muscle synergies, principal components, or continuum robotics may serve to speed up this bottleneck. In addition, the current implementation does not take full advantage of performance enhancements available in OpenSim and Simbody. Software optimizations could provide further performance enhancements. Another limitation is that the tracking error in rotational tasks is currently calculated using the Euler angle values, which are prone to well-documented numerical issues. In the future, spherical linear interpolation of quaternion rotations can be implemented for more robust error calculations in orientation tracking tasks. The current implementation is also currently available to users only through code (either C++ or SWIG-generated bindings) or OpenSim’s XML file format. Adding an interactive widget to the OpenSim GUI in the future will make the tool more easily accessible.

TSC provides a number of exciting opportunities for advances in simulation-based insights into neural control of gait and other movements. For example, TSC could be used to investigate high-level features, such as whole-body angular momentum or COM control, which are invariant across individuals and can be used to produce realistic gait patterns for a range of anthropometries. Because it is a predictive framework, TSC can also be used to simulate the effects of muscle fatigue or load carriage on gait patterns. TSC could also be used in conjunction with neural networks to perform physics-informed machine learning.

## Conclusion

Task space control is a paradigm widely used in robotics which has the potential to lead to new discoveries in neural control of human movement. This paper presents a mathematical overview of an implementation of floating-base task space control for biomechanics applications. The framework has been implemented in OpenSim in order to facilitate more widespread use among researchers in biomechanics. Example results from a gait simulation demonstrate the potential of the task space control framework to aid researchers in exploring neural control of movement.

## Acknowledgments

The authors would like to acknowledge Dr. Dimitar Stanev, whose initial implementation of a task space control framework in OpenSim formed the basis for this work. The authors also acknowledge Dr. Jeff Reinbolt for useful discussions on task space control, and Dr. Gary Zientara for his support of this effort. The authors also acknowledge Dr. Paulien Roos for management of the project under which this work was performed, and Ryan Middle and Garrett Tuer for their assistance in testing and reviewing the code. This material is based upon work supported by the U.S. Army Research Institute of Environmental Medicine under Contract No. W81XWH-22-C-0020. Any opinions, findings and conclusions or recommendations expressed in this material are those of the author(s) and do not necessarily reflect the views of the U.S. Army Research Institute of Environmental Medicine.

